# A Naturalistic Study on the Combined Neural and Psychological Effects of Psilocybin and Compassion Focused Imagery

**DOI:** 10.64898/2025.12.17.694940

**Authors:** Carla Pallavicini, Lorena Llobenes, Federico Cavanna, Laura Alethia de la Fuente, Stephanie Muller, María Costa, Natali Gumiy, Nicolás Bruno, Tomás D’Amelio, Jaskaran Basran, Ptarmigan Plowright, Ain Stolkiner, Mauro Namías, Paul Gilbert, Enzo Tagliazucchi

## Abstract

Psilocybin is a classic psychedelic drug known to alter subjective experience and elicit long-term psychological changes, enhancing cognitive flexibility and reducing rigid self-related beliefs. Combined with compassion motivational primes that involve generating mental representations of compassion, it may increase the potential for activating the care-affiliative motivational systems, linked to several important biopsychosocial processes underpinning social safeness, social connection and mental wellbeing. We investigated the synergetic effects of psilocybin and compassion imagery with self-reported questionnaires and functional resonance imaging data (fMRI) in a sample of 105 participants. Participants were primed with either attention to breathing or a short compassion focused imagery prime. We found a long-term synergetic effect of compassion imagery and psilocybin on cognitive absorption, as well as changes relative to baseline self-compassion and decentering. Based on functional interactions between attentional, executive and default mode networks, fMRI-based classifiers detected participant engagement in compassion focused imagery before psilocybin intake and distinguished compassion imagery vs. attention to breathing priming only the high dose of psilocybin. Our results support the potential for synergistic effects from combinations of psilocybin and compassion-based interventions to induce long-term psychological changes, reshaping the functional organization of large-scale brain networks. Future confirmatory studies of our exploratory analyses should be conducted to determine whether the combination of psilocybin and compassion-based practices promotes increases in caring and contemplative abilities, enhanced psychological flexibility and well-being.

## Introduction

Psychedelic substances, once shrouded in stigma and controversy, have recently garnered renewed attention for their potential therapeutic benefits (Griffiths et al., 2016; Gründer & Jungaberle, 2021; Kim et al., 2025; Nichols & Walter, 2001; Perkins et al., 2021). Among these substances, psilocybin, found in certain species of mushrooms, has emerged as a promising candidate for mental health intervention (Griffiths et al., 2016; Goldberg et al., 2020; Carhart-Harris et al., 2021; Goodwin et al., 2022). Recent research has revealed compelling evidence of psilocybin’s capacity to promote psychological well-being and induce enduring positive changes in personality traits (Bouso et al., 2015; Bouso et al., 2018; Griffiths et al., 2011; MacLean et al., 2011; Nichols, 2016; Nichols et al., 2017; Perkins et al., 2021). These long-term modifications can be elicited by a single dose of psilocybin and are closely related to the quality of the subjective experience during the acute psychedelic state (Garcia-Romeu et al., 2014; Griffiths et al., 2008, 2011, and 2016; MacLean et al., 2011; Millière et al., 2018). When generating altered states of consciousness, psilocybin can create peak experiences, also referred to as “mystical experiences”. These states are characterized by ego dissolution, forms of self-transcendence and inter-connectedness, feeling part of ‘all things’, oceanic boundlessness, and activation of positive affect sometimes referred to as awe and bliss (Griffiths et al., 2008; Tagliazucchi et al., 2016; Millière, 2017; Pallavicini et al., 2021). These changes have been linked to enduring changes in personality, behavior, and well-being (Griffiths et al., 2011). Moreover, neuroimaging studies have demonstrated that psilocybin may facilitate the activation of the core areas of what has been called the ‘social brain’ (Slavich et al, 2023) that evolved to facilitate attachment, social connectedness, friendship and cooperation (Rodríguez & Winkelman, 2021). These brain areas are particularly sensitive to social information in regard to the similarity, support and helpfulness of others and impact social cognition, and social reward systems, potentially offering insights into the therapeutic mechanisms of psilocybin (Carhart-Harris et al., 2012; Preller & Vollenweider, 2018; Nardou et al., 2023). In addition, compassion and compassion training have also been shown to impact many areas of the social brain, for example the ventral striatum, the (subgenual) anterior cingulate cortex, and the orbitofrontal cortex, along with neural networks associated with positive emotion and emotion regulation (Förster & Kanske, 2022; Kim et al., 2020; Vrtička, et al., 2017).

The Relaxed Beliefs Under pSychedelics (REBUS) model (Carhart-Harris & Friston, 2019) provides a theoretical account of the mechanisms through which psychedelics produce positive therapeutic outcomes. Grounded in the Bayesian predictive processing framework (Clark, 2013; Friston, 2010), REBUS proposes that psychedelics relax the precision-weighting of high-level priors (how strongly the brain relies on its top-down beliefs when interpreting reality), reducing the rigidity of maladaptive beliefs and increasing both neural and psychological flexibility. As with other facets of the psychedelic acute effects, this outcome is intricately tied to the context of the experience, also known as “set and setting”, the former referring to the brain state at the time of ingestion in regards to a sense of social safeness or unfamiliarity and threat, and the latter to the external environment (Hartogsohn, 2016; Studerus et al., 2012; Haijen et al., 2018; Carhart-Harris et al., 2018). While implementing settings conducive to the therapeutic action of psychedelics can be relatively straightforward, identifying and eliciting a favorable mindset remains more challenging, but in view of the processes involved in the evolution of the social brain (Slavich et al., 2023; Rodríguez & Winkelman, 2021) and their psychophysiological mediators, it is likely to be linked to issues of social trust and care-protection. Herein lies the potential value of contemplative practices as primes because these practices stimulate core brain areas linked to feelings of social connectedness and caring (Förster, & Kanske, 2022; Kim et al., 2020; Vrtička, et al., 2017). These conditions have already shown promise in eliciting positive outcomes when combined with psilocybin and other classic serotonergic psychedelics (Griffiths et al., 2018; Millière et al., 2018; Heuschkel et al., 2020; Holas et al., 2023; Timmermann et al., 2023).

Compassion-Focused Therapy (CFT; Gilbert, 2000, 2014, 2020 2022, 2023; Petrocchi et al., 2024) and Compassionate Mind Training (CMT) were developed to help individuals with high shame and self-criticism, who often respond poorly to traditional cognitive-behavioral interventions because of difficulties in generating self-soothing compassionate and supportive inner dialogue. Grounded in evolutionary psychology, attachment theory, and social mentality theory, CFT is a biopsychosocial therapy model that conceptualizes compassion as an evolved (stimulus-response guided) motivation, rooted in evolved caregiving systems. The evolution of caring helps the ‘cared-for’ to regulate threat and seek social connectedness, affiliation and friendships as major evolved biosocial goals (Rodríguez & Winkelman, 2021). Compassion, defined as a “sensitivity to suffering and needs in self and others, with a commitment to alleviate and prevent it” (Gilbert, 2014; 2022) therefore involves two parts: the stimulus which is the sensitivity to suffering and the response which is commitment to alleviating and preventing it. This motivation can operate through three flows: toward others, from others, and toward oneself; each supported by distinct psychophysiological mechanisms. These affiliative processes engage a balancing of the autonomic system with appropriate parasympathetic regulation via the vagus nerve, promoting states of safeness, affiliation, social trust, and flexible attention (Geller & Porges, 2014; Petrocchi & Cheli, 2019). Meta-analytic evidence across 47 studies indicates that CFT produces robust reductions in overall mental health difficulties across clinical and nonclinical populations, supporting its transdiagnostic efficacy (Petrocchi et al., 2024).

From a predictive processing perspective, both compassion training and psychedelics can be understood as processes that facilitate the updating of internal models (Clark, 2013; Friston, 2010; Carhart-Harris & Friston, 2019; Gilbert, 2020). Compassion reorients motivational systems from threat-sensitive, competitive strategies toward caring, prosocial motivations (Gilbert, 2014; 2022), thereby guiding the brain to update predictive models toward greater safeness and social connectedness. Psychedelics, by temporarily increasing cognitive flexibility and reducing the precision of rigid self-related beliefs, may potentiate this process (Carhart-Harris & Friston, 2019). Together, these approaches may create a synergistic dynamic in which insight and affiliation become mutually reinforcing, loosening rigid self-structures while anchoring expanded awareness in caring intention (Millière et al., 2018; Timmermann et al., 2023).

To empirically test this hypothesis, we designed a naturalistic field study combining psilocybin intake with compassion priming. Participants were randomly assigned to engage in either a short compassionate imagery or an attention-to-breath priming before the psilocybin session. Compassionate imagery was used to activate caring-based motivational systems through the generation of affiliative and soothing emotions, while attention to breath served as a structurally similar contemplative mindfulness task engaging attentional networks, as previously differentiated in the ReSource Project (Singer et al., 2016). The effects of psilocybin were controlled by comparing a high dose (HD) versus low dose (LD) experience, within an experimental protocol centered on the naturalistic use of psychedelics, avoiding the constraining factors often present in laboratory or clinical settings. The experimental design assessed subjective effects, changes in self-reported scales, and incorporated functional magnetic resonance imaging (fMRI) to elucidate the neural correlates of the observed effects, assessing whether changes in brain functional connectivity occurred synergistically due to the combination of compassion imagery priming and the high dose of psilocybin. Our main hypothesis was that psilocybin and compassionate imagery would interact synergistically on psychological processes underlying subjective well-being, with concomitant alterations in the functional connectivity of large-scale brain networks.

## Methods

### Overview of the experiment

The experimental procedure is illustrated in Fig. 1. Participants were recruited and screened according to the procedures outlined in the “Participants” subsection. After assessment of the inclusion criteria, a personalized timeline was established around each participant’s preferred intake day. One week prior to this day, participants completed questionnaires online and received Fitbit wristbands to record physical activity data, which they wore until one week after the experience. During the week preceding the intake day, participants visited the medical facility “Centro Diagnóstico Nuclear” (CDN) for fMRI scans (monitored for 5 minutes resting-state and 5 minutes compassionate imagery) and provided blood and saliva samples. Primary outcomes focused on contemplative abilities, including compassion and absorption. Secondary measures, such as personality traits and additional psychological variables, were collected but will be analyzed in future work.

**Figure 1.**
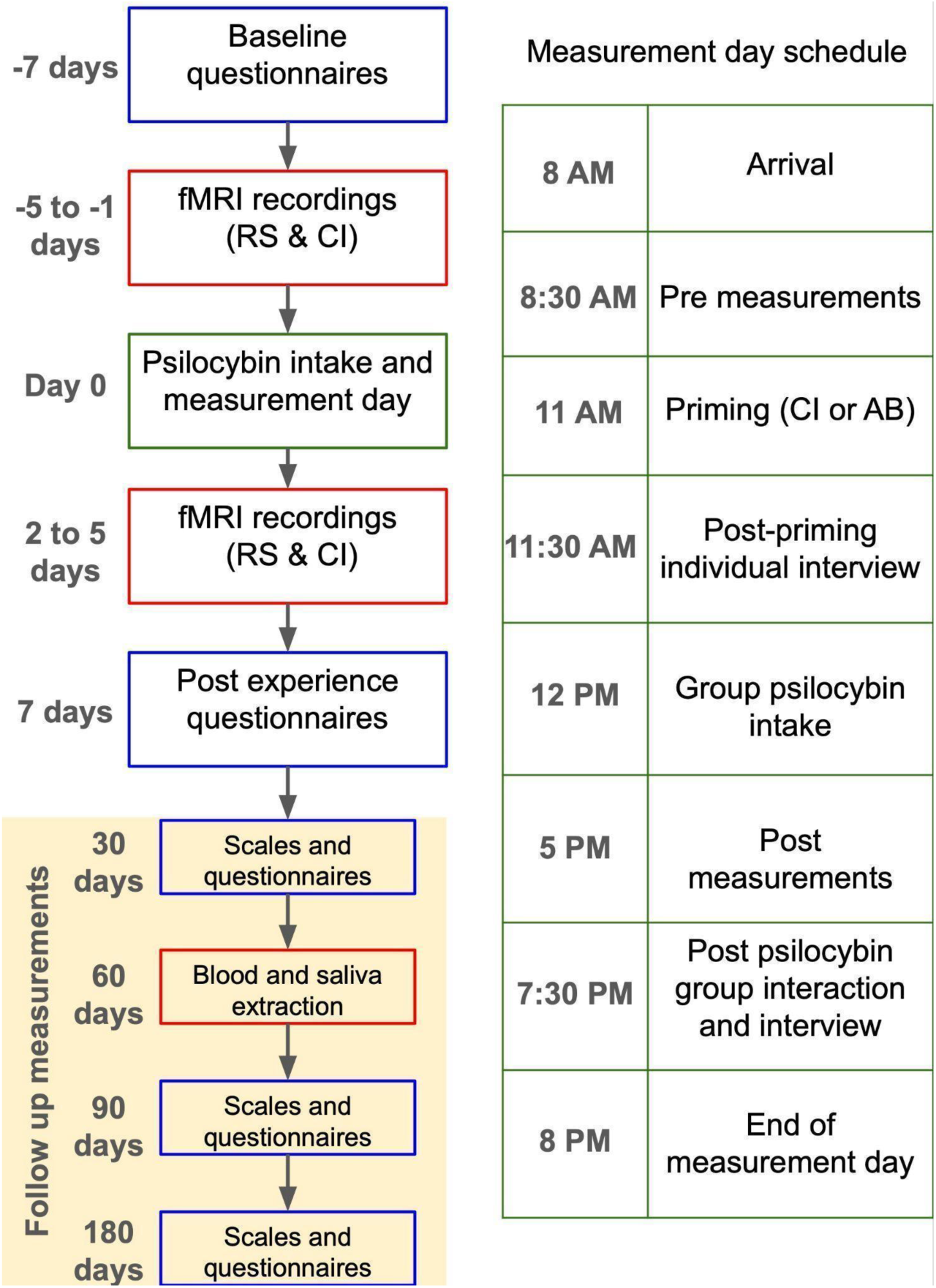
Overview of the experimental protocol. The left flow chart summarizes the questionnaires and biological/neuroimaging measurements performed before and after the psilocybin intake (day 0). The color of the boxes indicate the location of the measurements: remote (blue); medical center CDN (red), experience site (green). Neuroimaging acquisitions consisted of fMRI data acquired during resting state (RS) and during the practice of compassionate imagery (CI). Blood and saliva samples were obtained for analysis in a future report. The right flow chart presents the typical schedule of the psilocybin intake and measurement day. Before the group psilocybin intake, participants were briefly primed with either compassionate imagery or attention to breathing (AB).

On the day of the psilocybin intake, groups of 5 to 10 participants arrived at the location at 8 am to complete pre-experience assessments, including scales, questionnaires, interviews, and EEG recordings (5 minutes resting-state (RS) and 5 minutes compassionate imagery (CI)). Following this, participants underwent contemplative priming: Compassionate Imagery for active groups and Attention to Breathing (AB) for controls, lasting approximately 20 minutes. Individual interviews were then recorded. Subsequently, participants and their facilitator moved to a private area, with the time of psilocybin intake recorded. During the following 5 hours there was no intervention or interaction with researchers unless requested by a participant. After this period, participants underwent post-experience measures, including saliva extraction, scales, questionnaires, EEG recordings (5 minutes resting-state and 5 minutes compassionate imagery), and individual and group interviews. Participants left the location around 8 pm.

During the week after the ceremony, participants returned to the medical facility for follow-up MRI scans and blood and saliva samples. Fitbit bands were returned, and participants completed questionnaires online 7 days after the ceremony. Follow-up measures included online questionnaires at 1, 3, and 6 months after the intake day, with a blood draw at 2 months post-experience.

### Participants

Participants were recruited through word of mouth and social media advertising from October 2021 to June 2022. A screening process conducted by mental health professionals, involving the Depression Anxiety and Stress Scale - 21 (DASS-21) (Antony et al., 1998; Lovibond & Lovibond, 1995) and a non-diagnostic interview was used to assess volunteers’ suitability. During this initial screening, participants were briefed on the experiment’s details, objectives, and inclusion/exclusion criteria, which are detailed below.

Participants who fulfilled inclusion criteria received an information sheet outlining what the study included and an informed consent form. A total of 107 participants (52 female, 55 males ages 38 ± 8 years (mean ± std)) were recruited for the study. Two participants were unable to begin the experiment. As a result, 105 participants completed all assessments up to and including the psilocybin intake day. Five participants withdrew after the psilocybin intake day so 100 completed the post-intake fMRI session. Due to dropout, this reduced to 93 participants at follow up.

Participants had previously agreed with a facilitator to partake in a group experience with psilocybin and agreed to participate in the experimental protocol undergoing the measurements described in the previous subsection, which required blinding regarding the received dose of psilocybin. They were also informed that psilocybin would not be provided by the research team, and that the researchers would only conduct measurements before and after the acute effects.

The study was conducted in accordance with the Helsinki Declaration and approved by the Committee for Research Ethics at Universidad Abierta Interamericana University (Buenos Aires, Argentina) on May 31, 2021 (Reference Number: 0-1068).

### Inclusion and exclusion criteria

To be included in the experiment, participants were required to have a previous arrangement with the facilitator with the intention of participating in a group psilocybin experience. They also agreed to be blinded regarding the received dose of psilocybin. Participants agreed to receive contemplative training (CI or AB), undergo non-diagnostic EEG and fMRI scans, a non-diagnostic interview with a mental health professional, blood and saliva extractions and to wear a FitBit wristband during two weeks. Participants included in the experiment also agreed to avoid consuming psychoactive drugs at least one week before the beginning of the experiment, and to abstain from tobacco, alcohol and caffeine consumption at least 24 hours before their intake date.

All participants were aged between 21 and 65 years. Pregnant women were excluded from the experiment, as well as people with metallic material implanted in their bodies, people with pacemakers and claustrophobia. Participants who declared past difficult experiences with psychedelics with lasting negative psychological sequelae or who put themselves or others at risk were excluded from the experiment. A non-diagnostic psychiatric interview (SCID-CT; First, 2014) was conducted according to the guidelines by Johnson et al. (2008). Participants who fulfilled DSM-IV criteria for the following disorders were excluded from the experiment: schizophrenia or other psychotic disorders, and type 1 or 2 bipolar disorder (both also in first and second degree relatives), substance abuse or dependence over the last 5 years (excluding nicotine), depressive disorders, recurrent depressive episodes, obsessive-compulsive disorder, generalized anxiety disorder, dysthymia, panic disorder, bulimia or anorexia, as well as Participants with history of neurological disorders. Participants who presented one standard deviation above the mean in the State-Trait Anxiety Inventory (Spielberger, 2010) were excluded, as well as participants under psychiatric medication of any kind. All participants were required to be willing and able to provide informed consent.

### Experimental conditions and psilocybin administration

Once participants signed the informed consent, they were put in touch with the organizing team who received information concerning the planned psilocybin intake day. Participants chose their preferred date, and groups were arranged with the facilitator based on these preferences. As a control for psilocybin consumption and compassionate imagery priming, we used an active placebo consisting of a lower dose combined with a contemplative practice which only focused attention to breathing. The 100 participants who completed the post-measurements, were part of the following groups: High dose of psilocybin with CI priming (HD CI) (N = 30); High dose with AB priming (HD AB) (N = 20); Low dose with CI priming (LD CI) (N = 25); Low dose with AB priming (LD AB) (N = 25). The number of participants who completed follow up measures reduced to 93 and were part of the following groups: High dose of psilocybin with CI priming (HD CI) (N = 26); High dose with AB priming (HD AB) (N = 19); Low dose with CI priming (LD CI) (N = 23); Low dose with AB priming (LD AB) (N = 25). High doses comprised 3 gr of dry *Psilocybe* mushrooms and low doses comprised 1 gr of the same material.

Psilocybin experiences occurred in a variety of settings, encompassing both indoor and outdoor environments. Outdoor settings, particularly those immersed in natural surroundings with grass, trees, and open spaces, were predominantly favored for these experiences. Facilitators administered the doses by grinding the dry mushrooms and mixing the resulting powder with fruit juice, serving one cup per participant. The amount of powder per cup was weighed to ensure all doses were equal. From the moment of ingestion until 5 hrs after, participants only interacted with facilitators and their teams, unless they required special attention by the professionals in the research team. Facilitators provided recorded and live music, aromatics, and intervened if deemed necessary by accompanying participants during difficult parts of the experience.

### Psychometric, wellbeing and experience scales and questionnaires

Participants completed a series of scales and questionnaires at different time points during the experiment. One week before and after psilocybin intake, participants completed two sets of questionnaires online over 2 consecutive days. Set 1 included Spanish versions of various scales, such as the Big Five personality test (John and Srivastava, 1999), (STAI trait) (Spielberger, 2010), and Tellegen absorption scale (TAS) (Tellegen and Atkinson, 1974). Set 2 included additional scales, focusing on measures of compassion and contemplative abilities such as Compassion Engagement and Action Scales (CEAS) measuring compassion to others, from others and for self (Gilbert et al. 2017), and the Experiences Questionnaire (EQ), Decentering subscale (Fresco et al., 2007), among others.

On the day of psilocybin intake, participants completed pre-experience and post-experience questionnaires, assessing their subjective states and experiences with the drug. The pre-experience set included the Spanish versions of: State Anxiety Inventory (STAI state) (Spielberger, 2010), and MAIA scale (Mehling et al., 2012). The post-experience set included the Spanish versions of: State Anxiety Inventory (STAI state) (Spielberger, 2010), 21 items presented in the form of a visual analogue scale (VAS) adapted from Carhart-Harris et al., (2016) to determine the intensity of the acute effects, AWE (Yaden et al., 2019), the mystical experience questionnaire (MEQ-30), MEQ Setting, MEQ Social factors, MEQ Fusion (Barrett et al., 2015), 5D altered states of consciousness scale (5D-ASC) (Studerus et al, 2010) and MAIA (Mehling et al., 2012).

Follow-up measures were conducted virtually at 30, 60, and 180 days after the ceremony, including the same sets of questionnaires (Set 1 & 2) with the addition of an Events of Life Scale (Casullo 2004). Due to follow-up measure dropouts, the self-reported questionnaire dataset was reduced to 93 participants.

The complete list of measures are provided in Supplementary Material. Each questionnaire was scored as described in the corresponding citation. Those that presented unbalanced experimental grouping at baseline were excluded from the analysis.

### fMRI data acquisition

Two BOLD-weighted fMRI scans were acquired 2 to 4 days before and after the psilocybin intake: resting state (RS) and Compassionate Imagery (CI). The scanner model was Philips Ingenia, with a 3T field strength and a 16-channel phased array head coil with direct digital sampling. Both were obtained using a GRE-EPI imaging sequence, TR/TE = 3000/30ms, field-of-view = 240mm, 80 × 80 acquisition matrix, parallel acceleration factor (sense) = 3, 90° flip angle. Fifty-eight oblique axial slices were acquired in an interleaved fashion, each 3.0 mm thick with zero slice gap (3.0 mm isotropic voxels). The duration of each of the two BOLD scans was 5:00 minutes. For RS measurements, participants were instructed to stay still and awake in the scanner with no explicit instructions regarding the content of their thoughts. For the CI scan, a brief compassionate imagery instruction was given to the participants, who were asked to construct a compassionate image in their minds. fMRI data acquisition initiated immediately after the instruction was given.

### fMRI pre-processing

FSL tools were used to extract and average the BOLD signals from all voxels for each participant in each brain state. The preprocessing included 5 mm FWHM Gaussian spatial convolution, bandpass filtering between 0.01 and 0.1 Hz, and brain extraction (BET), followed by a transformation to a standard space (2 mm MNI brain). The next preprocessing steps were implemented in MATLAB, using in-house developed scripts. First, we corrected the data by performing regressions between the displacement parameters, the average signals extracted from the white matter and ventricles, their first derivatives, and the voxel-wise BOLD signals, retaining the residuals for further analysis. In the second step, we applied volume censoring (scrubbing) and discarded participants who presented significant relative head displacements in more than 20% of the recorded frames, with a criterion for movement significance set as a displacement between consecutive frames exceeding 0.5 mm (Power et al. 2014). Finally, we averaged all voxels within each ROI defined in the automated anatomical labeling (AAL) atlas, considering only the 90 cortical and subcortical non-cerebellar brain regions to obtain one BOLD signal per ROIs (Tzourio-Mazoyer et al. 2002). For RSN analysis, we averaged the signals corresponding to eight RSNs, the standard masks of the eight resting state networks (RSNs) as described in Beckmann et al. (2005) are freely available and were downloaded from http://www.fmrib.ox.ac.uk/analysis/royalsoc8/.

### Multivariate machine learning classifiers

We trained random forest classifiers (Breiman, 2001) to distinguish different types of fMRI measures (before vs after, RS vs CI, and all experimental group pairs) based on empirical individual FC matrices (fully connected weighted matrices computed using Pearson’s linear correlation coefficient between BOLD time series from each participant), and also based on RSN connectivity (RSNC), using a five-fold cross-validation procedure to estimate classifier accuracy. Random forest classifiers were trained using scikit-learn (https://scikit-learn.org/) (Abraham et al. 2014). We trained random forest classifiers with 1000 decision trees and a random subset of features of size equal to the rounded square root of the total number of features. The quality of each split in the decision trees was measured using Gini impurity, and the individual trees were expanded until all leaves were pure (i.e. no maximum depth was introduced). No minimum impurity decrease was enforced at each split, and no minimum number of samples was required at the leaf nodes of the decision trees (the classifier hyperparameters can be found in https://scikit-learn.org/).

To assess the statistical significance of the classifier accuracy values, we trained and evaluated a total of 1000 random forest classifiers using the same features (i.e. FC or RSNC matrices) as inputs but scrambling the class labels. We then constructed an empirical p-value by counting the number of times the accuracy of classifiers trained on scrambled class labels exceeded that of the original classifier. Classification performance was quantified using the area under the receiver operating characteristic curve (AUC), and results were considered significant at *p* < 0.05. The same permutation-based approach was applied to assess feature importance, comparing values from real versus scrambled-label classifiers.

### Statistical Analysis

Statistical analyses was conducted using the *statsmodels* Python library (https://www.statsmodels.org/stable/index.html). Normality was assessed using skewness and kurtosis values, which were all within the recommended parameters of 2 for skewness and 7 for kurtosis (West et al., 1996). Participants with significant missing data for measures at multiple timepoints were excluded from relevant analyses. Non-paired t-tests were used to compare self-reported questionnaire scores between groups and within groups between time points, determining statistical significance at p<0.05, FDR corrections were performed and differences sustained after correction reported. Correlational analyses between fMRI features and the scores of questionnaires were performed using Pearson’s linear correlation coefficient and reported significant at p<0.05 and at least a medium effect size for the correlation coefficient (|R|>0.3).

## Results

### Experimental protocol overview

The schedule for the measurements conducted before, during and after the psilocybin intake day are summarized in Fig. 1. One week prior to the psilocybin intake day, participants completed questionnaires online. During the preceding week, they also underwent fMRI scans which consisted of 5 minutes of rest alternated with 5 minutes performing a compassionate imagery task. On the day of psilocybin intake, groups of 5 to 10 participants arrived at the location at 8 am and completed a series of assessments, including questionnaires and recorded interviews. Following these measurements, the participants underwent priming with either short Compassionate Imagery (CI) or Attention to Breathing (AB), each lasting approximately 20 minutes, followed by another round of individual interviews. During the following 5 hours, a facilitator administered psilocybin as previously arranged with the group of participants, without intervention nor interaction with the researchers. Subsequently, participants completed questionnaires to assess the experience. During the following days, the subjects took part in another fMRI recording session with identical instructions and parameters as the one completed before dosing. Participants also completed scales online 7, 30, 90 and 180 days after the experiment. Besides the measurements summarized in Fig. 1, we also acquired other data sources (e.g. actigraphy, pre and post dosing EEG, blood and saliva samples) that will be analyzed and reported in future publications.

### Acute effects during the psychedelic experience

The radar plots in Fig. 2 show the outcome of questionnaires related to the subjective experience elicited by psilocybin, including subscales from the MEQ (Mystical Experience Questionnaire) (Barrett et al., 2015), VAS (Visual Analogue Scales) (adapted from Carhart-Harris et al., 2016), 5D Altered States of Consciousness (5D-ASC) (Studerus et al., 2010), and Awe (Yaden et al., 2019). The left panel compares HD vs. LD, while the right panel compares HD CI vs. HD AB. In the HD vs. LD comparison, significant differences (uncorrected) were observed in four dimensions, including MEQ Transcendence, 5D audiovisual synesthesia, 5D anxiety, and 5D complex imagery. However, these accounted for only a small subset of the 20 dimensions tested, indicating that the two doses elicited broadly similar acute subjective experiences. No significant differences were found between the HD CI and HD AB groups, suggesting that the low-dose active control and the priming conditions were effective in minimizing potential unblinding due to subjective effects.

**Figure 2.**
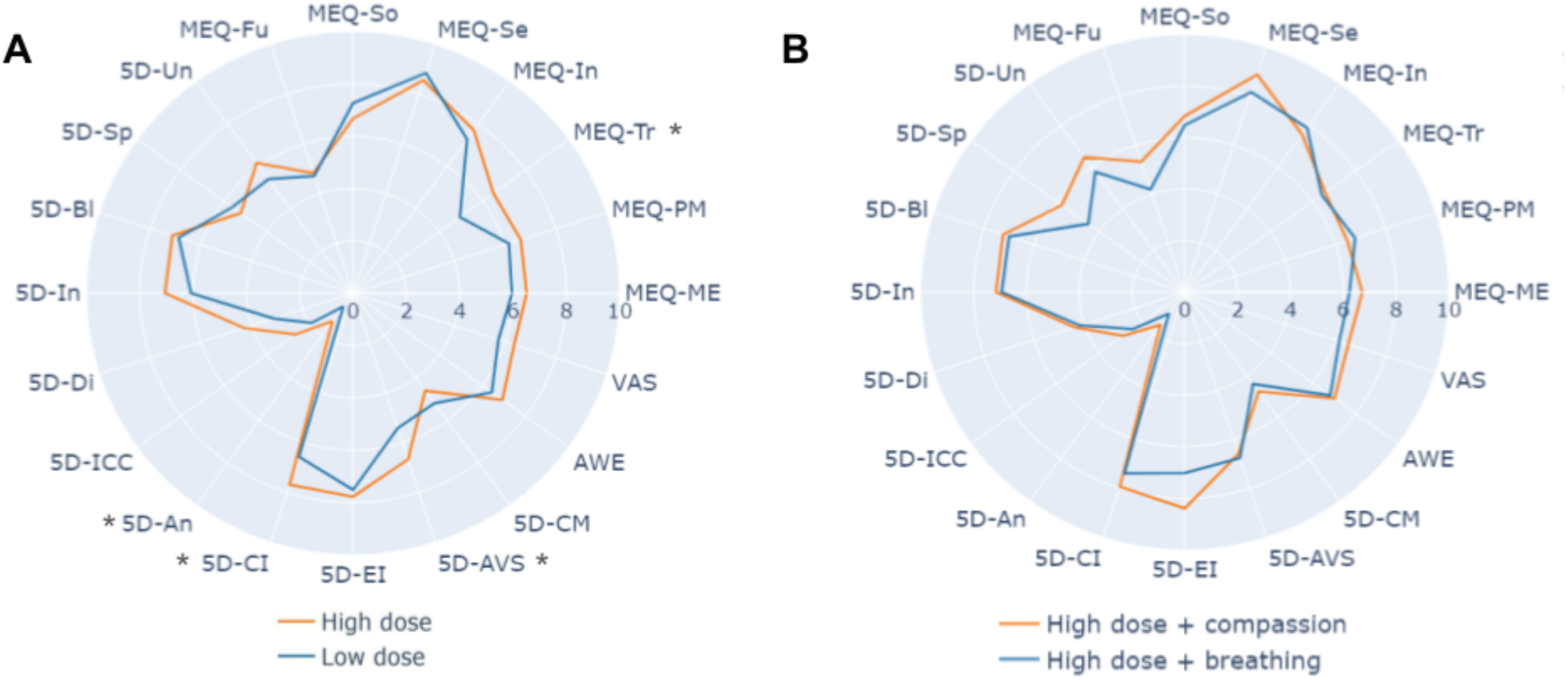
The low and high dose conditions resulted in similar profiles of subjective effects. The radar plot on the left summarizes the questionnaires obtained after high and low doses, including the following scales and subscales: MEQ (Mystical Experience Questionnaire), VAS (Visual Analog Scale), 5D (5D Altered States of Consciousness Questionnaire) and Awe. The abbreviations corresponding to each item are described in Supplementary Table 1. The radar plot on the right presents the same information but compared between compassion imagery and attention to breathing priming, in both cases for the high dose condition (* p<0.05).

### Effects of psilocybin and priming on questionnaires

Participants responded to a series of questionnaires 7 days before the psilocybin intake day, and 7, 30, 90 and 180 days after that event. For each questionnaire subscale, we performed the following comparisons: HD vs. LD, and HD CI vs. HD AB. The latter comparison aims to capture the synergistic effects of priming and a high dose of psilocybin. We performed comparisons between groups for each time point, as well as within-group comparisons between different time points. We excluded subscales (compassion towards others, CEASTowards, and from others CEASFrom), which presented significant differences at baseline from these analyses, indicating that the self-reports made by the different groups of participants were not matched for that dimension (see Supplementary Fig. S1). Also 2 of the 3 dimensions of the Compassion Fears scales were unbalanced at baseline which drove us to disregard this measure as well (Supplementary Fig. S2).

Figure 3 summarizes the results of comparing questionnaire subscales between conditions, with shaded gray background indicating p<0.05 between groups (FDR corrected), and solid and dashed brackets indicating increases from baseline to the time points FDR corrected and uncorrected, respectively. Uncorrected findings are reported as preliminary trends that may warrant further investigation in future studies. On the Tellegen Absorption Scale (TAS), participants in the HD condition showed significant FDR-corrected increases relative to baseline at +30, +90, and +180 days. In contrast, the LD condition exhibited only a tendency towards increased absorption at +90 and +180 days, which did not survive correction for multiple comparisons. In the priming comparison, TAS scores were significantly higher in the HD CI group than in the HD AB group at +30, +90, and +180 days (all FDR-corrected). Within-group analyses further revealed that the HD CI group presented robust increases from baseline to these same time points (all FDR-corrected), while no significant longitudinal effects were observed for HD AB.

**Figure 3.**
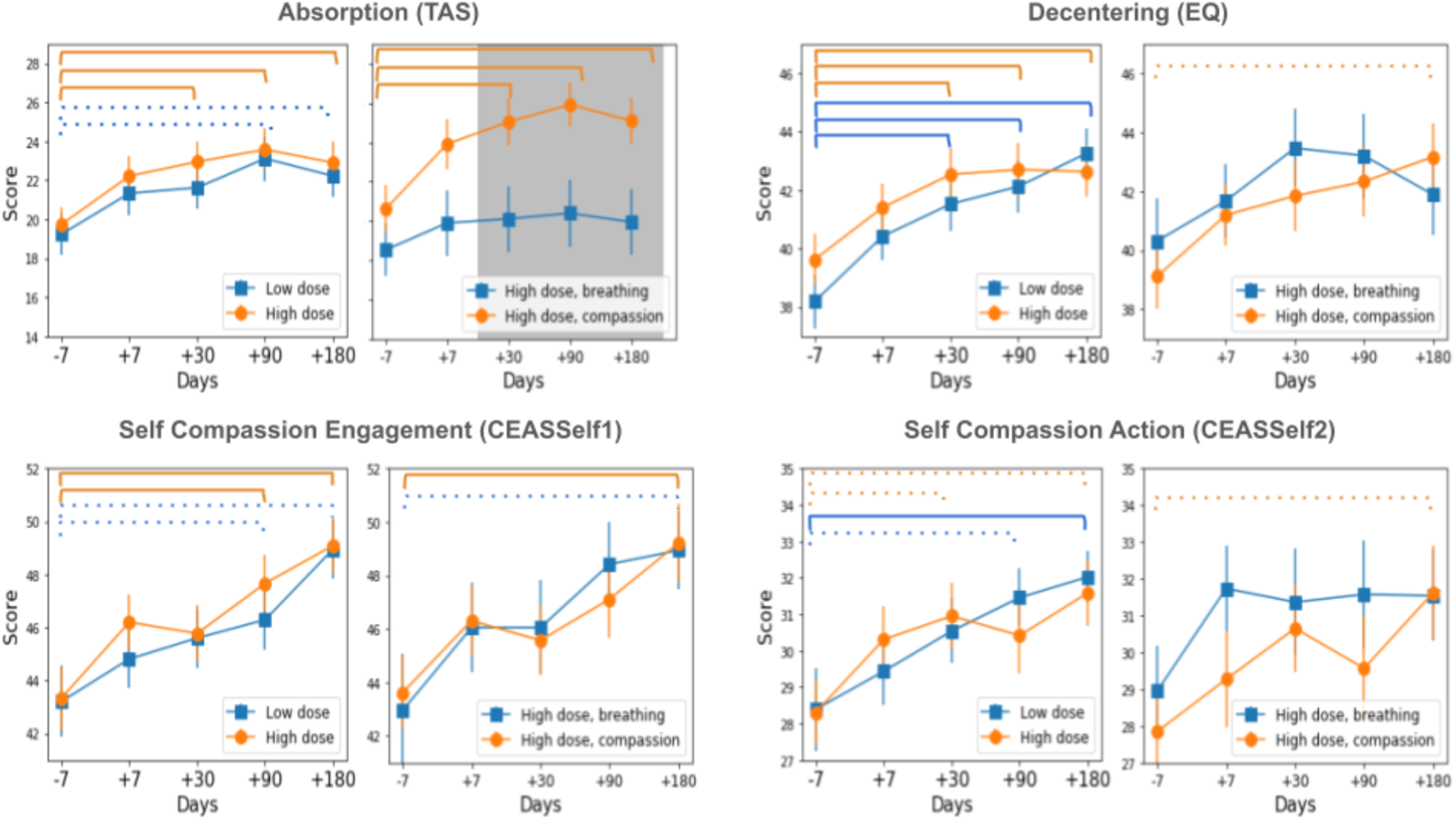
Psilocybin produced a synergistic effect with compassionate imagery on absorption and was associated with within-group increases in decentering and self-compassion. Each subplot summarizes results for low vs. high dose (left) and for compassionate imagery vs. attention to breathing priming (right) in the high dose. Results are shown for multiple time points before and after the psilocybin intake day. Shaded gray background indicates p<0.05 (FDR corrected) between groups. Orange and blue brackets indicate significant (p < 0.05) differences between the time points for the corresponding group color. Solid brackets indicate FDR correction passed.

On the Experiences Questionnaire (EQ) – Decentering subscale, both HD and LD conditions showed significant FDR-corrected increases at multiple time points after psilocybin intake. For the priming comparison, we observed a trend towards greater increases in the HD CI group relative to baseline, with the effect approaching significance at +180 days, whereas no such effect was observed for HD AB.

On the Self compassion scales (CEASSelf1 & CEASSelf2), we observed differential within-group effects for engagement and action over time. For CEASSelf1 (engagement), the HD group showed significant FDR-corrected increases relative to baseline at +90 and +180 days. The LD group displayed similar tendencies at the same time points, though these did not survive correction. Within the priming conditions, the HD CI group exhibited a significant increase at +180 days, while HD AB only showed a non-significant tendency. For CEASSelf2 (action), both HD and LD conditions displayed tendencies towards increases over time, with only the LD group at +180 days surviving FDR correction. In the priming comparison, the HD CI group presented a trend towards increased action at +180 days, though this effect did not reach FDR significance. It is important to note that these findings reflect within-group temporal changes and do not imply statistically significant between-group differences. While the observed patterns varied across groups, the current results do not provide sufficient evidence to conclude that one condition or intervention was more effective than another. Nevertheless, reporting these trends is valuable for generating hypotheses and identifying potentially meaningful directions for future research, particularly given the exploratory nature of the study and the novelty of combining psilocybin with compassion-based interventions.

Overall, these results indicate that psilocybin intake, particularly at high doses and when combined with compassionate imagery, was associated with sustained increases in absorption, and showed trends toward enhanced decentering and self-compassion engagement up to six months after the experience, warranting further investigation.

### fMRI-based classification of experimental conditions

Based on the result of previous neuroimaging studies, we assessed whether the experimental conditions had an impact on whole-brain functional connectivity (FC) by training and assessing a multivariate machine learning model for their classification. This model consisted of a random forest classifier trained to distinguish between baseline resting state (RS) and compassionate imagery functional connectivity (CI FC), and between changes in CI FC before and after the psilocybin intake, for subjects primed or not with compassionate visualization. The first of these two tasks aimed to detect differences intrinsic to the CI practice, while the second aims to classify CI FC scans based on the effect on the activity elicited by the CI priming to psilocybin. We evaluated classifier performance for both the HD and LD conditions and found significant classification only in the HD group (Supplementary Fig. S3).

Each machine learning classifier was trained using the FC between an established set of RSN comprising primary and extrastriate visual networks (VI and ES), an auditory network (AU), the sensorimotor cortex (SM), the default mode network (DM), the fronto-parietal executive control network (EC), and the right/left dorsal attention network (DR and DL) (Beckmann et al. 2005). For each evaluation, 1000 random forest classifiers were constructed, both with real data and scrambled labels for the calculation of p-values associated with the classifier accuracy and the feature importance reflecting the relevance of each pairwise FC for the classifier, as described in the Methods section. The accuracies were determined using the Area Under the Curve (AUC) of the Receiver Operating Characteristic (ROC) curve (<AUC>) of the 1000 repetitions.

As shown in Fig. 4, the classifier successfully distinguished between both sets of experimental conditions. In the case of RS vs. CI, this classification was predominantly based on increased FC values between the DR and DL, and between these two networks and EC, DM and AU (<AUC> = 0.716 +-0.008; p < 0.001) (Fig. 4A). In the comparison of connectivity differences before and after intake in the CI task (ΔCI) between the AB and CI priming, we recovered a significant positive classification (<AUC> = 0.696 ± 0.008; p < 0.05) for the high dose conditions (Fig. 4B). We found significant increases in the coupling between the DR and DL networks, and between AU and both SM and DR. However, the same comparison for the low dose groups did not result in accurate classification (<AUC> = 0.49 ± 0.01; p > 0.5), illustrated in Supplementary Figure 2.

**Figure 4.**
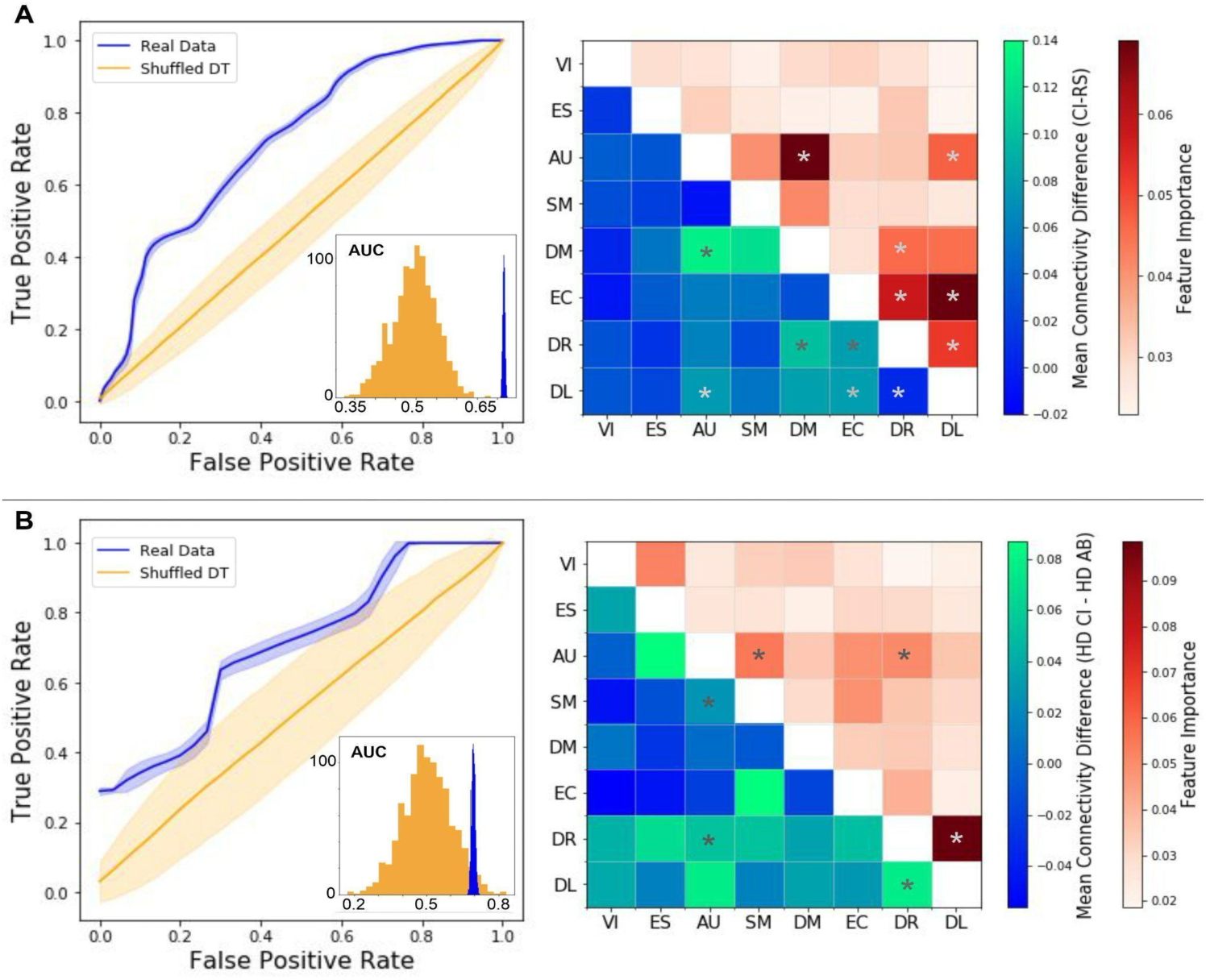
Whole-brain functional connectivity allows discriminating between RS and CI (A), and between CI vs. AB priming in the HD (B). The left panels show the ROC plot obtained using data with shuffled labels (orange) and with the unshuffled, original data (blue). Insets show the AUC distributions obtained after 1000 random permutations of the labels. The right panel presents matrices with the above diagonal entries representing the mean feature importances associated with each functional connection between pairs of RSN, and below diagonal entries indicating differences between groups (RS - CI, and HD CI - HD AB) (*p<0.05).

### Correlations between fMRI and self-reported questionnaires

Afterwards, we investigated the correlations between changes in RSN FC and the dimensions of the self-reported questionnaires (measured on the same time-point as the fMRI data) for which we previously found a significant effect of dose, or a synergistic effect of dose and priming (Fig. 3). The results of this analysis are shown in Fig. 5. Importantly, all correlations were computed on change scores in functional connectivity during the compassionate imagery (CI) task (ΔFC = POST CI - PRE CI) rather than on absolute FC measures. Thus, the associations below should be interpreted as linking individual variability in CI-evoked connectivity change to questionnaire scores.

**Figure 5.**
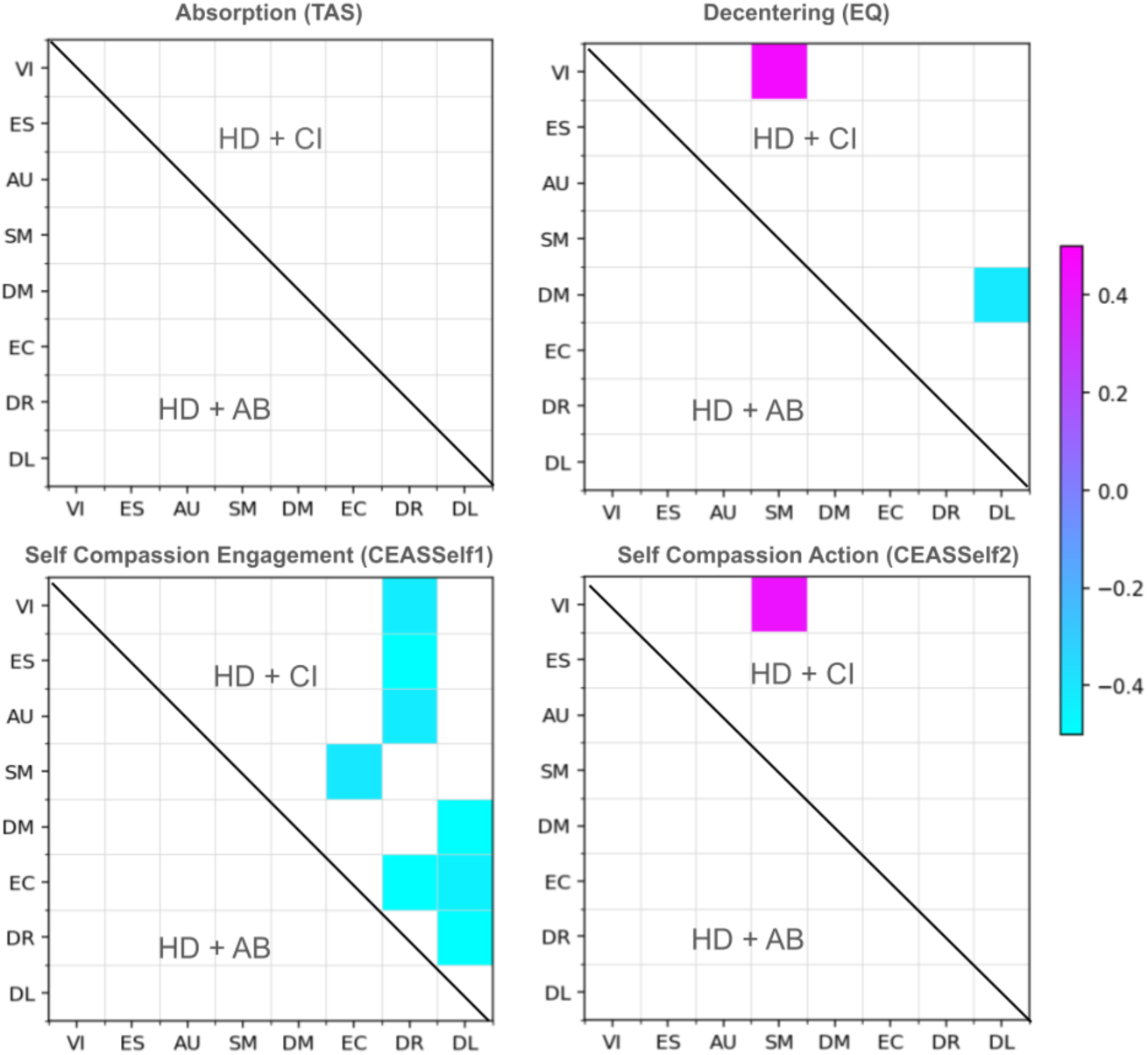
Changes in fMRI FC during compassionate imagery present mostly negative correlations with the dimensions affected by the psilocybin high dose, and by the combination of psilocybin and compassionate imagery. For each self-reported dimension, the matrix elements indicate the correlation with the changes in FC between the corresponding RSN (indicated in the x and y axes) for the HD CI (above diagonal) and HD AB (below diagonal). Only entries with |R|>0.3 and p<0.05 are shown in the figure.

TAS showed no significant correlations in either dose condition. By contrast, all other scales exhibited significant associations only in the HD CI group. Specifically, EQ (Decentering) and CEASSelf2 (action) showed positive correlations with connectivity between visual cortex and sensorimotor (VI–SM) networks, indicating that larger increases in decentering and self-compassionate action were associated with stronger VI–SM integration across sessions. In turn, CEASSelf1 (engagement) displayed predominantly negative correlations, including reduced coupling between the dorsal attention networks (DL, DR) and the rest of the RSNs, and a negative EC–SM association.

## Discussion

Although both psychedelic intake and compassionate imagery have been individually associated with increased psychological flexibility and adaptive emotional processing, their combined effects have not been previously explored. This study assessed the potential synergistic impact of psilocybin and compassion focused imagery on key markers of well-being, including decentering, compassion, and absorption, known to support emotional regulation and resilience. We employed self-report questionnaires assessing stable traits and psychological processes, alongside fMRI recordings acquired before and after psilocybin intake, followed by either compassionate imagery priming or a control priming based on attention to breathing. Given the novelty of this interaction, our approach was exploratory in nature.

Since the contemporary renewed interest in psychedelic-assisted psychotherapy, obtaining a reliable control for placebo effects has remained a persisting challenge, given the difficulties of masking the subjective aspects of the experience (Luoma et al., 2020; Nayak et al., 2023). To address this issue, we proposed an active placebo condition consisting of 1g of dried mushrooms, as opposed to the high dose of 3g. Additionally, consumption occurred in a naturalistic setting where groups of participants shared the experience. Interestingly, when comparing the self-reported subjective experience during the acute effects of high vs. low doses of psilocybin, we found only limited differences across a few dimensions, and no significant effects between HD CI and HD AB (Fig. 2). This result is indicative of an effective blinding of the control condition, considering that even though participants did not report significant differences in their subjective experiences, we could nevertheless differentiate those groups based on questionnaire and neuroimaging data. Our ability to successfully blind the placebo condition is a notable strength of our study, as it enhances the internal validity of our findings. However, this blinding may also contribute to smaller differences between the research groups. It is well-documented that outcomes in psychedelic research often have an expectancy component, where participants’ beliefs can influence the results (Olson et al., 2020; Muthukumaraswamy et al., 2021; Aday et al., 2022; Szigeti et al., 2024). In our study, if a substantial proportion of participants believed they were receiving the high dose, this belief could have attenuated the differences between the experimental and control groups, potentially leading to smaller observed effect sizes.

An interesting result reflected in our data is the consistent increase in absorption, as measured by the Tellegen Absorption Scale (TAS). The HD group showed robust, FDR-corrected increases at multiple follow-up time points, while the LD group displayed non-significant tendencies in the same direction. This suggests that psilocybin generally enhances absorption in the experience, but higher doses are needed for reliable, sustained effects. Absorption describes the propensity to become fully engaged in perceptual or imaginative experiences (Tellegen & Atkinson, 1974) and has been linked to openness to altered states of consciousness and to contemplative practices such as meditation (Pekala et al., 1985; Radtke & Stam, 1991; Cuevas-Vargas et al., 2023; Yang et al., 2024). In this sense, increased absorption may be viewed as an externally driven contemplative ability (Lifshitz et al., 2019), with the psychedelic state potentiating immersion in inner imagery or sensory experience (Aday et al., 2021). We also observed a synergistic effect: the HD CI group showed significantly greater increases in TAS relative to HD AB, both at specific time points and across baseline-to-follow-up comparisons. This suggests that compassionate imagery not only shaped the acute psychedelic state but also potentiated a longer-lasting enhancement in absorption. Such outcomes resonate with evidence linking contemplative training to absorption-related capacities (Millière et al., 2018) and highlight the potential of combining psychedelics with structured practices to consolidate contemplative traits. In this context, enhanced absorption may underlie the sustained ability to engage deeply with compassionate imagery and other self-transcendent experiences, bridging acute psychedelic phenomenology and enduring changes in well-being (Haijen et al., 2018; Lifshitz et al., 2019; Aday et al., 2021).

Another finding was the consistent increase in decentering, as measured by the Experiences Questionnaire (EQ). Both HD and LD groups showed significant FDR-corrected increases at multiple follow-up time points. A trend toward stronger effects was observed in the HD CI group at six months, indicating that compassionate imagery may modestly potentiate this trait over time. Decentering refers to the metacognitive ability to observe one’s thoughts, feelings, and internal experiences as temporary, objective events in the mind rather than as accurate reflections of the self or reality (Fresco et al., 2007). It involves a shift from being immersed in one’s inner narrative to adopting a witnessing or observational stance, which enables greater psychological flexibility and emotional regulation. This ability supports adaptive self-reflection and has been linked to cognitive reappraisal, the process of changing the meaning of experiences by re-evaluating emotions toward them (Gross & John, 2003, Berenstein et al., 2015; Kobayashi et al., 2020). Recent work highlights decentering as a key mechanism in contemplative practices, including mindfulness-based interventions and compassion-based training, where it underpins resilience and psychological flexibility (Campo & Yali, 2025). Importantly, emerging evidence suggests that psilocybin may facilitate both self-compassion and decentering by disrupting rigid self-referential processing, loosening habitual patterns of cognitive fusion, and fostering openness to alternative perspectives and modes of self-relatedness (Diehl & Rosenthal, 2024). These shifts are thought to underpin therapeutic change by enhancing psychological flexibility, a core process targeted in various transdiagnostic interventions (Pilecki et al., 2024). Enhancing decentering creates a psychological stance conducive to flexibility, reduced rumination, and prosocial engagement. The trend toward stronger effects in the HD CI group suggests that compassionate priming may consolidate this process, extending the overlap between psychedelic phenomenology and compassionate imagery. By fostering the capacity to observe thoughts and emotions without over-identification, psilocybin combined with compassion imagery may provide a pathway toward lasting improvements in psychological flexibility and well-being.

Another key finding of this study was the increase in self-compassion, particularly in the high-dose condition, with trends toward stronger effects when combined with compassion imagery (Fig. 3). To our knowledge, this is the first report of a link between measures of compassion and psychedelic use. Self-compassion refers to the ability to direct the evolved motivation to care and alleviate suffering inward, fostering a courageous and wise stance toward one’s own pain (Gilbert 2000; 2014). It involves two interrelated components: *engagement,* the motivation to face and understand suffering in self and others rather than avoiding, denying or suppressing it, and *action*, the intentional effort to alleviate that suffering through helpful thoughts or behaviors (Gilbert 2020; 2022; Petrocchi et al 2024). Rooted in the evolved care motivational system that promotes caring and responses to being cared for, self-compassion activates physiological states of safeness and affiliation that counteract threat responses and regulation (Kim et al., 2020). Research has shown that cultivating self-compassion through CFT is associated with reduced self-criticism, shame, anxiety, and depression, as well as improved emotion regulation and resilience (Gilbert & Procter, 2006; Petrocchi et al., 2024). More recently, Petrocchi et al. (2024) found that CFT produced a large and consistent increase in self-compassion in both clinical and non-clinical populations. There is now growing evidence for an association between self-compassion and a range of health-promoting behaviors, including greater psychological flexibility and resilience (Homan & Sirois, 2025; Eghbali et al., 2022; Tiwari et al., 2020).

In summary of our self-report findings, we observed a robust, synergistic effect on absorption, with significantly greater increases in the high-dose compassionate imagery group compared to other conditions. While increases in decentering and self-compassion were also observed over time within certain groups, these did not yield significant between-group differences and should be interpreted as trends. Together, these mechanisms may establish the conditions necessary for the care-affiliative motivational systems to be selectively engaged and sustained over time, providing a plausible neuropsychological account of sustained effects observed up to six months following psilocybin administration.

Our analysis of fMRI recordings successfully differentiated between resting state and compassionate imagery tasks performed within the scanner (Fig. 4A), highlighting the recruitment of different neural mechanisms with each task. The classification of these two brain states primarily relied on functional connectivity (FC) among the left and right dorsal attention networks (DL and DR), as well as between these networks and the fronto-parietal executive control network (EC), default mode network (DM), and auditory network (AU). The dorsal attention networks (DANs) encompass the dorsal frontal and parietal cortices, specifically the intraparietal sulcus (Beckmann et al., 2005). These networks are implicated in the voluntary orientation and maintenance of attention to a location (Corbetta et al., 2000) and are considered goal-driven attentional networks that utilize internal goals or expectations to focus on sensory stimuli (Corbetta et al., 2008). There is evidence linking the connectivity of these networks with mindfulness and contemplation traits (Parkinson et al., 2019; Froeliger et al., 2012), connecting the functions of the DANs to the contemplative requirements of sustained, focused attention. Hence, we can relate these facets of contemplative practices to compassionate imagery and support the conclusion that this practice is also related to increases in DANs inter and intra-connectivity, associated with a more focused state of mind. Another key observation is that nearly all resting-state network connectivities contributing to the distinction between RS and CI were higher during the imagery condition. Since resting-state networks typically emerge from patterns of mutual anti-correlation between distinct functional systems, this widespread increase in connectivity suggests that compassionate imagery evokes a more integrated and globally connected brain state. Such enhanced network integration parallels previous reports of increased global connectivity during psychedelic states, hinting that both conditions may share a neural signature of broadened integration and flexible information exchange.

Whole-brain functional connectivity also allowed the distinction between the two forms of meditation priming (AB vs. CI) in the high dose condition (Fig. 4B), suggesting that psychedelics enhance the recruitment of neural activity implicated in compassionate imagery priming, which differs from the neural activity patterns elicited by the control priming. Once again, connectivity within both the DR and DL networks was crucial for this classification, along with connectivity between the auditory network (AU) and both the sensorimotor network (SM) and DR. The involvement of the SM network may reflect the embodiment experience promoted by the CI task, as sensorimotor integration is central to grounding the bodily self and linking perception to action (Shimada et al., 2022; Ferri et al., 2012). Moreover, all significantly discriminative connectivity features were positive, suggesting that the HD CI group exhibited larger changes in connectivity compared to HD AB, reflecting a more pronounced neural impact of compassionate imagery under psychedelic influence. Given that we observed connectivity changes pre- and post-intake, these findings indicate a synergistic effect of psilocybin and CI on connectivity associated with attention orientation and sustainment, and embodiment. Considering available evidence linking regions of the insula to compassionate states and traits (Kim et al. 2020, Novak et al 2022), and taking into account that this area is a component of the AU (Damoiseaux et al. 2006), we consider that our results involving the connectivity to this network point towards CI priming exerting a significant effect detectable in terms of brain activity as measured with fMRI.

Lastly, we explored the relationship between changes in brain functional connectivity during the compassion imagery task and self-reported measures, focusing on the comparison between the HD CI and HD AB groups to more directly assess the synergistic effects of psilocybin and compassion priming (Fig. 4). Notably, no significant correlations emerged in the HD AB group, suggesting that psilocybin combined with attention-to-breath priming may result in weaker or less consistent links between brain dynamics and self-reported psychological outcomes. In contrast, multiple significant correlations were observed in the HD CI group, highlighting a stronger and more targeted impact of compassionate imagery when combined with psilocybin. A particularly relevant finding concerns the SM, a key hub in task-driven processes and in grounding the inner narrative self (Ferri et al., 2012; Weiss, 2013). This area’s positive connectivity with EQ and CEASSelf2 suggests that enhanced integration of embodied and perceptual networks may support the capacity to disengage from rigid self-referential processes and promote compassionate action (Farb et al., 2007; Ferri et al., 2012). In parallel, negative correlations between the DL and the DM with decentering reinforce the idea that psychological flexibility depends on rebalancing attentional and self-referential systems (Farb et al., 2007; Christoff et al., 2016). Furthermore, CEASSelf1 was associated with widespread negative correlations involving both dorsal attention networks (DR and DL), suggesting that reduced connectivity between these networks and the rest of the brain may facilitate a more focused, inwardly directed mode of compassionate engagement (Corbetta et al., 2008; Parkinson et al., 2019; Garrison et al., 2015). These results resonate with the CFT model, which distinguishes two complementary components of self-compassion: engagement and action. The engagement aspect (CEASSelf1), characterized by reduced connectivity across dorsal attention and executive–sensorimotor networks, reflects a shift away from externally oriented, goal-directed processing toward a more open and receptive internal awareness. In contrast, the action component (CEASSelf2) was linked to increased visual–sensorimotor connectivity, indicative of enhanced embodiment and perception–action integration, which may underlie compassionate responding. Together, these findings highlight the central role of task-positive networks (SM, DR, DL) and their interaction with self-referential systems in mediating the psychological effects of psilocybin combined with compassion imagery. The alignment of these neural signatures with reported gains in decentering and self-compassion highlights a potential mechanism through which contemplative and caring capacities are cultivated at the network level. The consistent implication of the dorsal attention networks further suggests a robust and specific role for these systems in supporting the mental states fostered by compassion imagery, warranting further exploration in future research.

The results of our study present a series of limitations. While investigating psychedelics in a natural context presents multiple advantages, it also imposes limitations related to lack of control in the administered doses, and to the difficulty of adopting and following a double-blind design. We could distinguish between low and high psilocybin doses; however, each dose was administered to a different group of participants, thus preventing us from performing within-group comparisons. This limitation was manifest in some metrics that presented significant differences at baseline, indicating that groups were not perfectly matched in that dimension. Moreover, the lack of significant results in the acute effects measurements could stem from participants lacking an adequate reference value to judge the intensity of the effects. Due to time constraints, the training in compassion-focused imagery received by the participants was brief at 20 minutes, therefore future studies with higher intensity or longer duration compassion-focused training are needed to replicate the findings. Finally, the non-invasive nature of fMRI recordings should be complemented with the analysis of biological markers, which will be reported in future publications.

In conclusion, our study shows that the combination of psilocybin and compassion-focused imagery produces sustained increases in absorption, and suggests trends toward enhanced decentering and self-compassion, psychological processes that may contribute to greater flexibility and well-being. Absorption provides the attentional immersion necessary to engage with inner experience, decentering allows individuals to relate more flexibly to thoughts and emotions, and compassion activates the caring system, creating a sense of safeness. Together, these mechanisms appear to interact synergistically: compassion imagery, particularly in high-dose psilocybin contexts, appears to function as an effective priming mechanism that may enhance the therapeutic potential of psychedelic experiences. By activating affiliative motivation and soothing physiological systems, compassion-based interventions may deepen key experiential processes such as absorption, decentering, and self-compassion, while promoting greater neural and psychological integration. The specificity of the effects observed in the high-dose compassion imagery (HD CI) group highlights the relevance of emotional safeness and affiliative states as modulators of psychedelic experiences. Accordingly, future research could investigate the integrative potential of Compassion-Focused Therapy (CFT) and Compassionate Mind Training (CMT), both as preparatory practices and post-experience frameworks to support sustainable therapeutic and self-transformative outcomes.

## Acknowledgements

This study was supported by funding from Spinoza (Spinoza Company B.V.) and The Compassionate Mind Foundation, Derby, UK. This project was also funded by grants ANID/FONDECYT Exploración 13240170 and ANID/FONDECYT Regular 1220995 (Chile)

## Supplementary material

**Figure S1:**
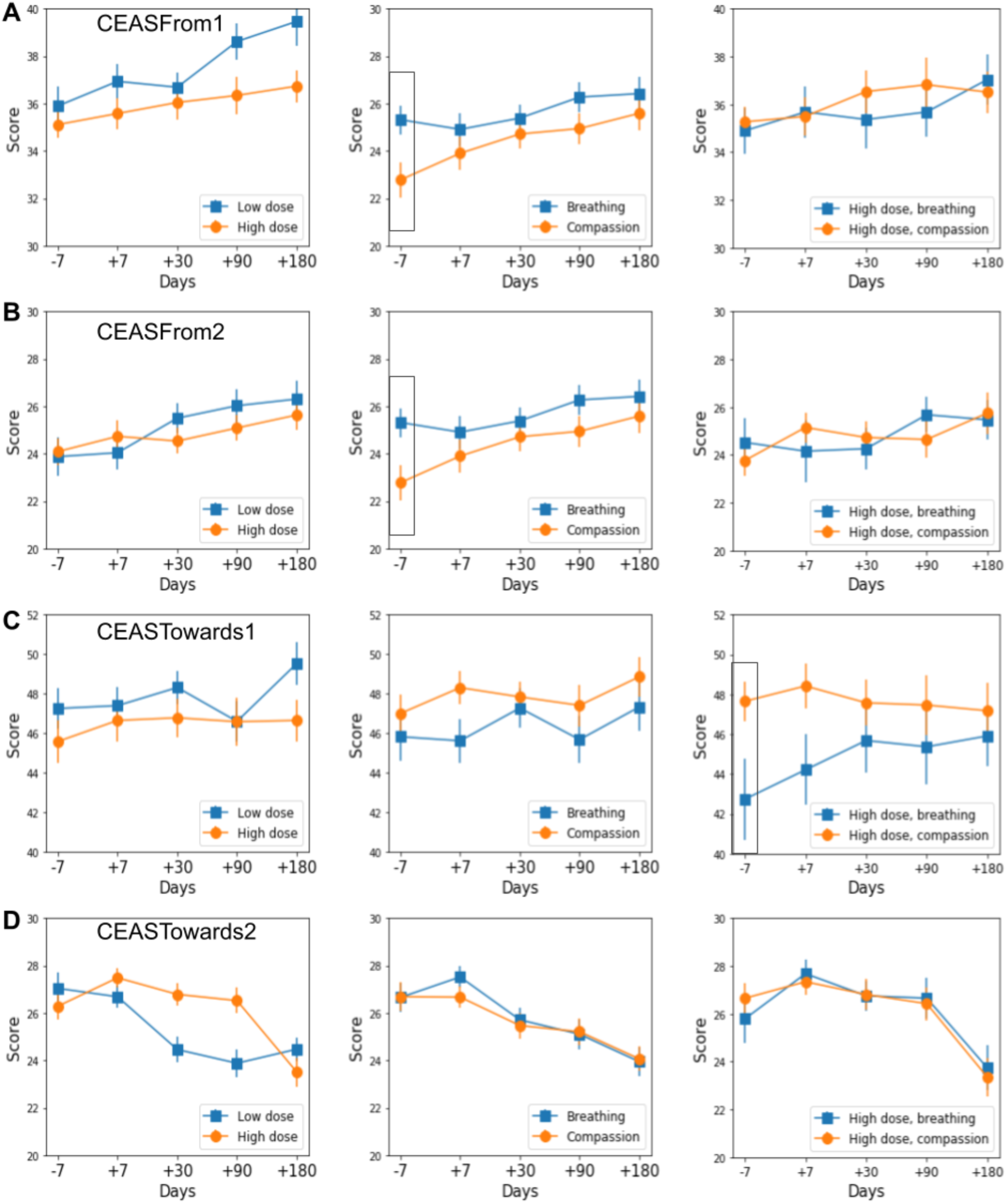
CEAS flows unbalanced at baseline were excluded from further analysis to avoid bias in interpretation. Each row represents a different CEAS subscale: CEASFrom1 (A), CEASFrom2 (B), CEASTowards1 (C), and CEASTowards2 (D). The column plots display comparisons for HD vs. LD (left), CI vs. AB (middle), and HD CI vs. HD AB (right). Although only the CEASTowards1 (engagement) subscale showed a significant baseline difference, we opted to exclude the entire flow from group comparisons to maintain analytic consistency and avoid selective reporting. Brackets highlight unbalanced baselines.

**Figure S2.**
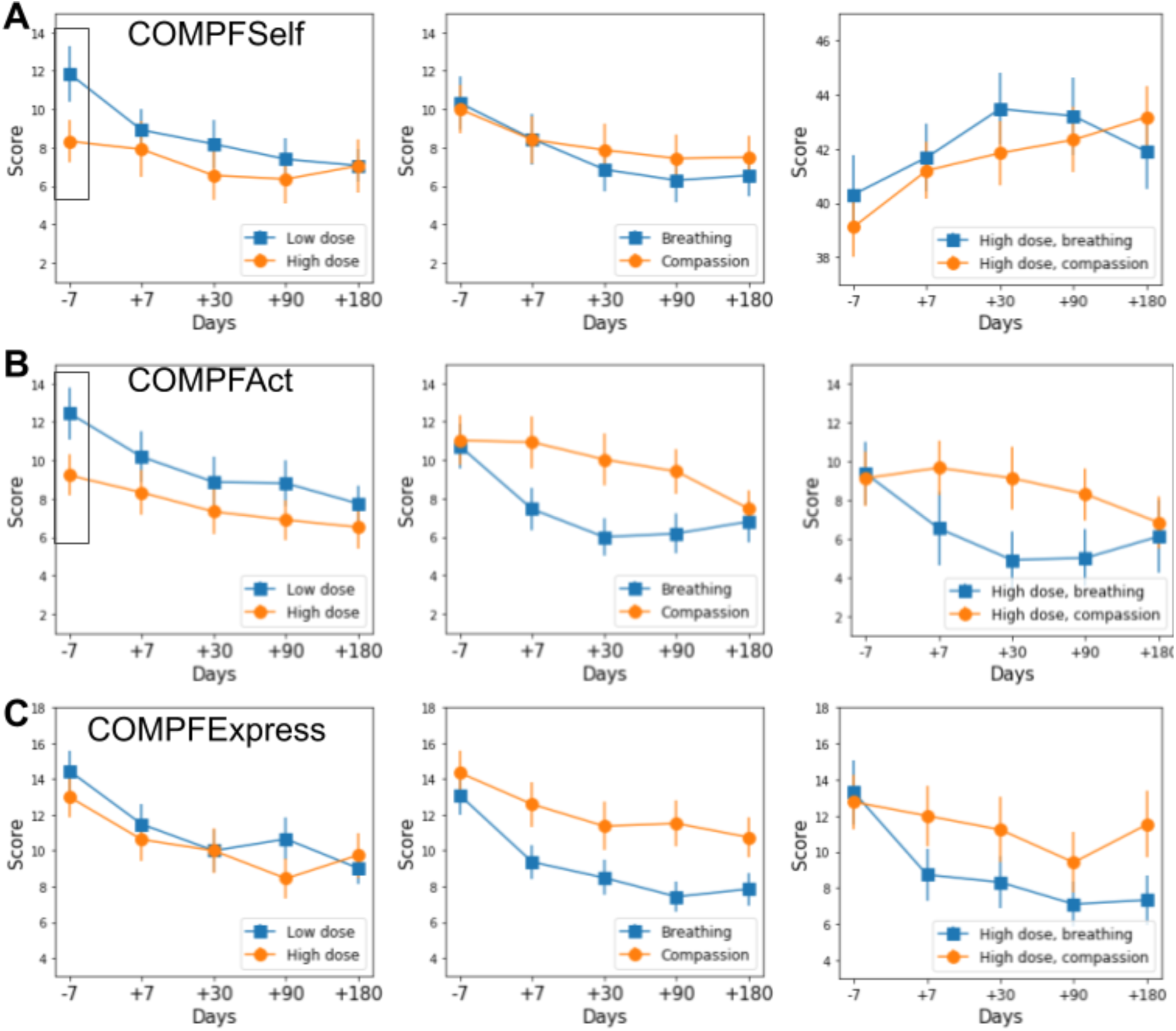
Unbalanced baselines in fears of compassion subscales. Each row displays a different subscale of the Fears of Compassion scale: COMPFSelf (A), COMPFact (B), and COMPExpress (C). Columns show group comparisons across time for Low vs. High dose (left), Breathing vs. Compassion priming (middle), and High Dose CI vs. High Dose AB (right). Timepoints span from baseline (–7) to +180 days. Two of the three subscales showed significant baseline differences across groups, which led us to exclude this entire metric from further group-level comparisons to avoid biased interpretation and selective reporting. Baseline imbalances are indicated with brackets.

**Figure S3.**
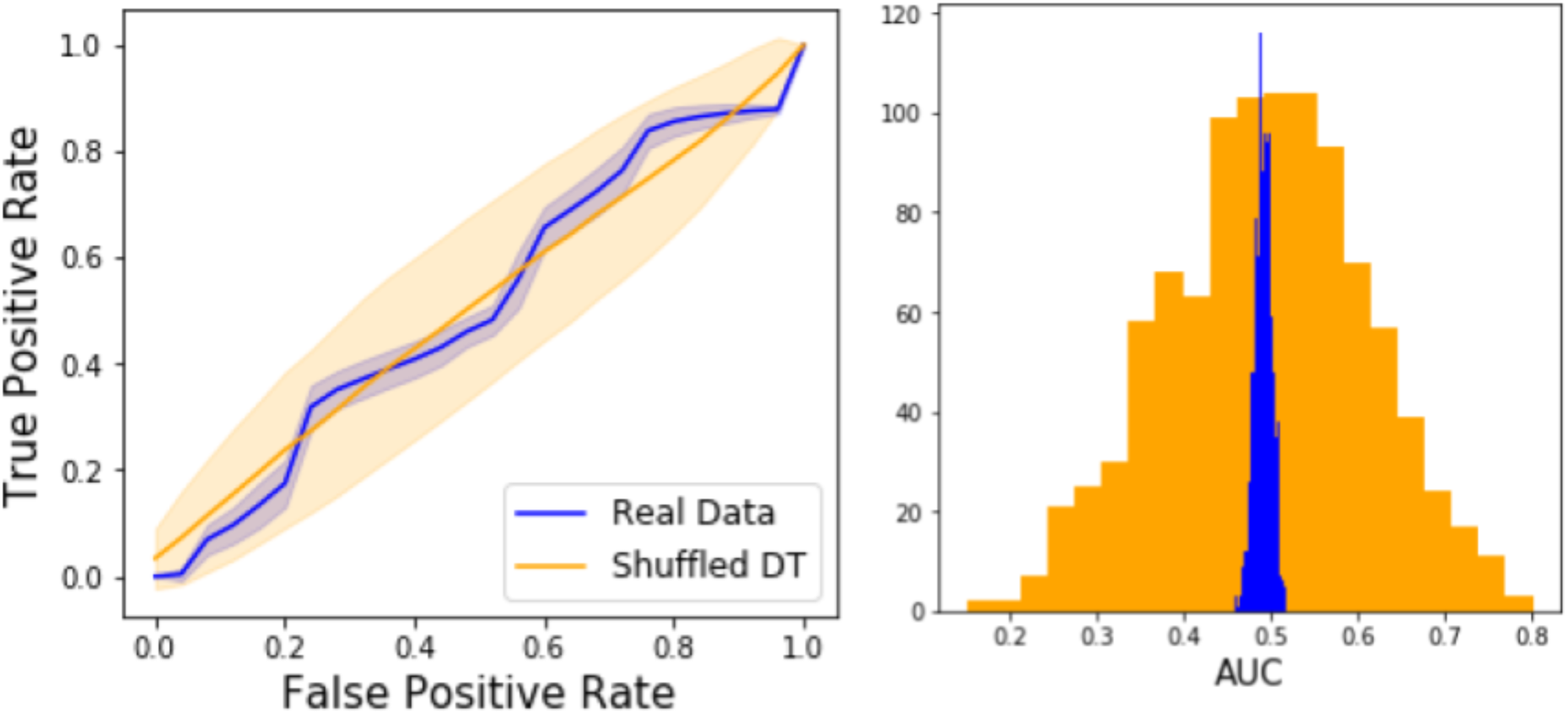
Classification of functional connectivity changes for the low dose condition between the Compassionate Imagery (CI) and Attention to Breath (AB) priming conditions. The left panel displays the ROC curve using the original data (blue) and the distribution under label permutation (orange). The right panel shows the distribution of AUC values across 1000 random label permutations. Classification performance did not exceed chance level (mean AUC = 0.49 ± 0.01, p = 0.516), indicating the absence of a reliable distinction between the CI and AB groups in the low dose condition.

**Supplementary Table 1.** List of all scales and questionnaires corresponding to each set. First Column presents the acronym, second the complete name of the questionnaire and the third lists all the subscales and dimensions within it.

| **Set 1** |  |  |
| --- | --- | --- |
| *BFI* | Big Five Inventory | Extraversion, Agreeableness, Conscientiousness, Neuroticism, Openness. |
| *STAI* | State-Trait Anxiety Inventory | Trait anxiety. |
| *PANAS* | Positive And Negative Affect State | Positive affect, Negative affect. |
| *TAS* | Tellegen Absorption Scale | Total score. |
| *BIEPS* | Escala de Bienestar Psicológico | Acceptance and control, Autonomy, Ties, Projects. |
| **Set 2** |  |  |
| *CEAS* | Compassionate Engagement and Action Scales | To self 1, To self 2, Towards others 1, Towards others 2, From others 1, From others 2. |
| *COMPF* | Compassion Fears | Express, Act, Self. |
| *SPS* | Social Safeness and Pleasure Scale | Total score. |
| *EQ* | Experiences Questionnaire | Decentering |
| *SCS* | Self-Compassion Scale | Total score. |
| *CNS* | Connectedness to Nature | Total score. |
| **Pre-experience Set** |  |  |
| *STAI* | State-Trait Anxiety Inventory | State anxiety. |
| *MAIA* | Multidimensional Assessment of Interoceptive Awareness | Noticing, Not-distracting, Not-worrying, Attention regulation, Emotional awareness, Self-regulation, Body listening, Trusting. |
| **Post-experience Set** |  |  |
| *STAI* | State-Trait Anxiety Inventory | State anxiety. |
| *VAS* | Visual Analogue Scale | Total score (VAS). |
| *AWE-S* | Awe Experience Scale | Time, Self-loss, Connectedness, Vastness, Physiological, Accommodation. Total Score (AWE) |
| *MEQ* | Mystical Experience Questionnaire | Mystical experience (MEQ-ME), Positive mood (MEQ-PM), Transcendence (MEQ-Tr), Ineffability (MEQ-In), Setting (MEQ-Se), Social (MEQ-So), Fusion (MEQ-Fu). |
| *5D-ASC* | Five Dimensional Altered States of Consciousness | Unity (5D-Un), Spiritual (5D-Sp), Blissful (5D-BI), Insightfulness (5D-In), Disembodiment (5D-Di), Impaired consciousness and cognition (5D-ICC), Anxiety (5D-An), Complex imagery (5D-CI), Elementary imagery (5D-EI), Audio-visual synesthesia (5D-AVS), Changed meaning (5D-CM). |
| *MAIA* | Multidimensional Assessment of Interoceptive Awareness | Noticing, Not-distracting, Not-worrying, Attention regulation, Emotional awareness, Self-regulation, Body listening, Trusting. |

